# A non-living, microbe-based surface coating promotes tubeworm and coral settlement

**DOI:** 10.64898/2026.08.06.743049

**Authors:** M.V. Farrell, L. Rix, P.A. O’Brien, T.L. Dunbar, S. Mahesh, F. Kuek, N.J Shikuma

**Affiliations:** Department of Biology and Viral Information Institute, San Diego State University, San Diego, California 92182 USA; School of Chemistry and Molecular Biosciences, Australian Centre for Ecogenomics, The University of Queensland, Brisbane, Queensland, Australia; Australian Institute of Marine Science, Townsville, Queensland, Australia

**Author notes:** Co-first authors (M.V. Farrell, L. Rix, P.A. O’Brien). **Corresponding author:** Nicholas J. Shikuma.

## Abstract

A major barrier to scaling marine restoration and aquaculture is the lack of reliable tools to induce invertebrate larvae to settle and metamorphose when and where needed. Although microbial cues are known to induce metamorphosis in many invertebrates, existing methods rely on natural biofilms that are variable, difficult to standardize, and unsuitable for large-scale deployment. Here we introduce ReefTiles, a non-living bacterial coating that preserves inductive activity from metamorphosis-stimulating marine bacteria in a stable, reproducible format. Using both tubeworm and coral larvae, we show that dried and inactivated bacterial films retain full settlement-inducing capacity, matching or exceeding live biofilms while eliminating concerns associated with releasing viable microbes into the environment. Viability assays confirm inactivation, and the coating adheres reliably to common substrate materials. Because ReefTiles can be manufactured and stored at scale and tailored to different inductive strains, they provide a practical microbe–based biotechnology for enhancing larval settlement in reef restoration, sustainable aquaculture, and engineered marine infrastructure.

## INTRODUCTION

Across restoration, aquaculture, and marine infrastructure industries, a persistent challenge is the reliable induction of larval settlement and metamorphosis in marine invertebrates. The transition from a planktonic larva to a benthic juvenile represents a critical developmental and ecological bottleneck for many species—including corals, oysters, abalone, and tubeworms—because it determines both population recruitment and the establishment of sessile communities (Cavalcanti et al., 2020; Holbrook et al., 2018).

In restoration settings, the inability to predictably stimulate larval settlement limits efforts to reseed degraded reefs and benthic habitats (Turnlund et al., 2025). For corals in particular, restoration projects have traditionally relied on outplanting fragmented adults, which are genetically uniform and vulnerable to stressors such as heat, disease, and bleaching (Boström-Einarsson et al., 2020). Enabling sexually produced coral larvae to settle and metamorphose would increase the genetic diversity and adaptive potential of restored populations, but achieving consistent recruitment under laboratory or field conditions remains challenging (Banaszak et al., 2023; Randall et al., 2020).

A similar problem exists in commercial aquaculture, where larval metamorphosis is often a point of massive loss. Mortality during this stage can decimate oyster or abalone hatcheries and is exacerbated by the absence of reliable cues to promote healthy settlement and post-metamorphic growth (Joyce & Vogeler, 2018). In both aquaculture and restoration, current methods for encouraging settlement—such as conditioning surfaces in natural seawater—are time-consuming, unpredictable, and prone to contamination by pathogens or competitors (Banaszak et al., 2023; Randall et al., 2020). This lack of control over the settlement process represents a technological and biological bottleneck across multiple industries that depend on the successful recruitment of sessile marine animals.

Existing methods for promoting larval settlement in restoration and aquaculture rely largely on conditioning artificial substrates in natural seawater, where bacterial and algal biofilms gradually accumulate over weeks to months (Ritson-Williams et al., 2010). These biofilms can elicit settlement in some species but are highly variable in composition and efficacy, depending on environmental conditions such as temperature, light, water chemistry, and microbial community structure (Kegler et al., 2017; O’Brien et al., 2025; Padayhag et al., 2023). This variability often leads to inconsistent recruitment outcomes and introduces the risk of pathogens or nuisance organisms that can outcompete or infect target species (Banaszak et al., 2023). Because the conditioning process depends on uncontrolled natural inputs, it is inherently difficult to reproduce across locations or timeframes, limiting its scalability for industrial or restoration applications. Chemical inducers such as potassium chloride, epinephrine, or histamine have also been used to trigger metamorphosis in annelids, mollusks, and corals (Carpizo-Ituarte & Hadfield, 1998; Erwin & Szmant, 2010; Joyce & Vogeler, 2018). However, these compounds act non-specifically and can “short-circuit” normal developmental signaling pathways, resulting in incomplete or abnormal metamorphosis and poor post-settlement survival.

Many marine invertebrates rely on bacteria as natural cues to initiate metamorphosis and settlement. In coral reefs, bivalve hatcheries, and benthic ecosystems, larvae encounter specific microbial biofilms that provide chemical or structural signals interpreted as suitable habitat. Classic model systems such as the tubeworm *Hydroides elegans* have revealed that certain bacteria produce molecular cues that can precisely trigger larval metamorphosis (Cavalcanti et al., 2020; Malter et al., 2024; Shikuma et al., 2014). Similar bacterial associations have been reported for corals, where compounds such as tetrabromopyrrole, cycloprodigiosin, or cell surface polysaccharides promote settlement in otherwise inactive larvae (Alker, Farrell, Demko, et al., 2023; Fiegel et al., 2025; Sneed et al., 2014a). These findings suggest that larvae have evolved to recognize conserved microbial cues, integrating bacterial presence into developmental decision-making. Several studies have shown that monospecies or defined multispecies bacterial biofilms can induce larvae settlement, including strains isolated from crustose coralline algae (Negri et al., 2001; Petersen et al., 2021; Tran & Hadfield, 2011). However, maintaining these biofilms requires continuous culturing and tightly controlled conditions that limit their practicality for large-scale or field deployment.

Despite the growing understanding that bacteria influence animal development, these insights have not yet been translated into scalable or practical technologies. A central challenge is identifying bacterial taxa or their products that are effective and sufficiently stable for use outside controlled laboratory settings. Natural biofilms are dynamic, fragile, and inconsistent, and maintaining live bacteria on deployment surfaces introduces ecological and regulatory risks in aquaculture and open-water environments. These limitations underscore the need for approaches that preserve inductive capacity without relying on living biofilms, enabling the development of reproducible, shelf-stable materials that harness natural bacterial cues to direct larval settlement for restoration, aquaculture, and broader marine biotechnology applications.

## RESULTS

### Challenge for reliable induction of larval settlement of marine invertebrates

To address the challenge of inconsistent larval settlement across marine restoration and aquaculture systems, we first developed a conceptual framework illustrating where a bottleneck occurs in current practice (**Fig. 1A**). Although corals, oysters, tubeworms, and other benthic invertebrates routinely produce microscopic swimming larvae, few successfully attach and metamorphose under hatchery or restoration conditions. This inefficiency limits both the genetic diversity and scalability of outplanting efforts. To overcome this constraint, we designed ReefTiles—substrates coated with bacterial biomolecules that mimic natural microbial cues—to provide a non-living, reproducible and shelf-stable settlement surface (**Fig. 1B**). Visual inspection showed that *Phaeobacter gallaeciensis* (Ruiz-Ponte et al., 1998) bacterial coatings produced visible films on ceramic surfaces. Once dried or heated, ReefTiles maintained an even, matte coating, indicating firm bacterial adhesion and persistence of surface material (**Fig. 1C**). This approach bridges the gap between laboratory discovery of bacterial inducers and practical deployment tools.

**Fig 1.**
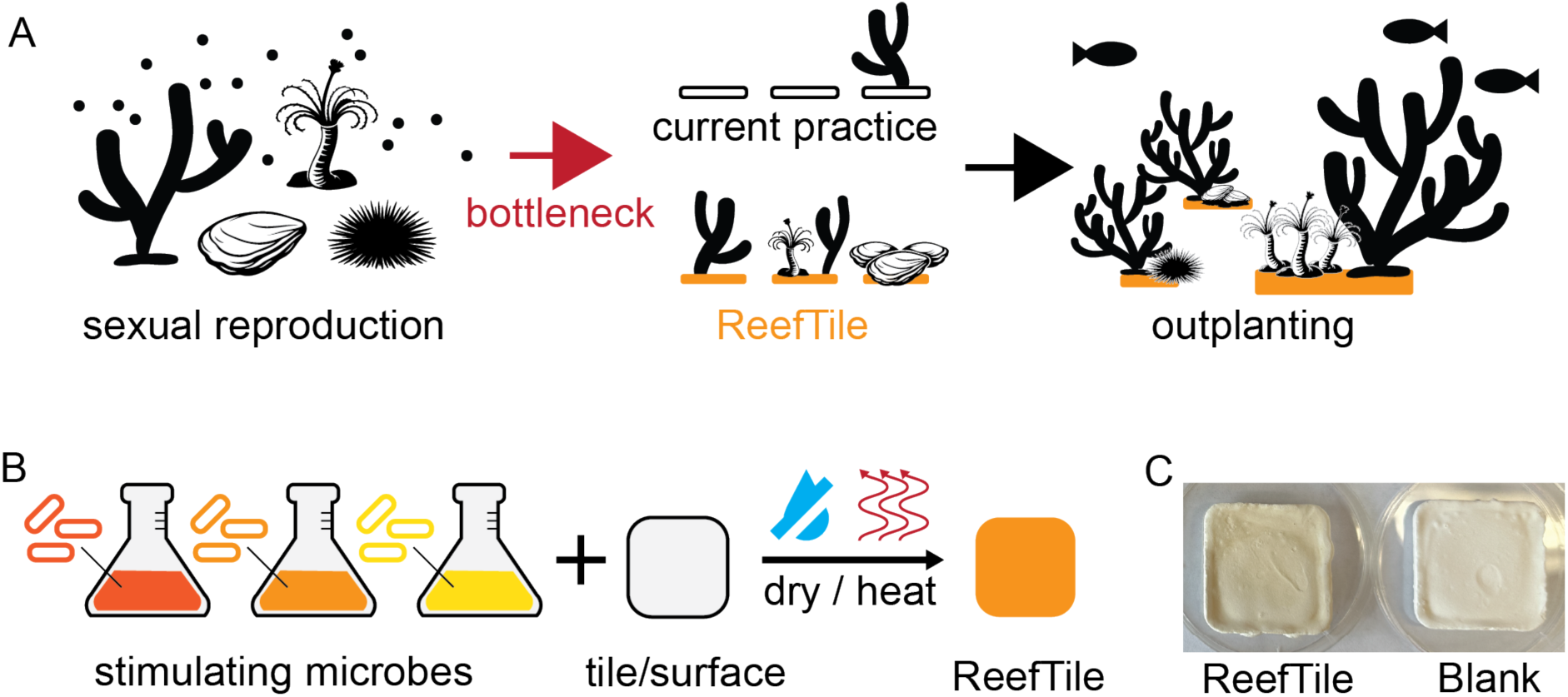
ReefTiles overcome larval settlement bottlenecks in marine restoration and aquaculture. **(A)** Many benthic marine animals—including corals, oysters, tubeworms, and sea urchins—reproduce sexually, releasing planktonic larvae that must locate a suitable surface for metamorphosis and attachment. In current practice, low and inconsistent larval settlement represents a major bottleneck to restoration and aquaculture success. ReefTiles, coated with microbial stimulants, provide a reproducible surface that promotes efficient settlement and growth, facilitating subsequent outplanting of diverse species to the natural environment. **(B)** Schematic of ReefTile preparation. Cultures of settlement-stimulating marine bacteria are grown and applied to ceramic or other substrates, then dried and/or heat-treated to inactivate cells while retaining stimulatory biomolecules. The resulting inert, bioactive coating yields a standardized ReefTile capable of inducing larval settlement across multiple marine taxa. **(C)** Image of ceramic settlement tile with visible ReefTile bacterial coating versus an unconditioned blank tile.

### ReefTile coating is inert and long-lasting

To test whether ReefTile coatings could retain stimulatory activity while remaining biologically inert, we compared freshly conditioned tiles containing live bacterial biofilms of *Phaeobacter gallaeciensis* with ReefTiles that had been dried over several days or heat-treated at 65°C. Live/dead fluorescence imaging confirmed that immediately after heat treatment, the *P. gallaeciensis* bacteria within the ReefTile coating were non-viable, in contrast to untreated biofilms that contained abundant live bacteria (**Fig. 2A**). We next investigated a panel of six bacterial strains previously associated with coral and tubeworm metamorphosis I ncluding strains isolated as part of the Reef Restoration and Adaptation Program(**Table S1**) (Rix et al., 2024). To quantify this loss of viability across all bacterial strains, bacterial colony counts were performed on freshly coated surfaces before and after the heating/drying process. For all six bacterial strains, no viable cells were detected following the ReefTiles treatments of heating or drying (**Fig. 2B**). Together, these results demonstrate that the coating procedure effectively inactivates bacteria while preserving a durable, adherent layer of bacterial biomolecules.

**Fig 2.**
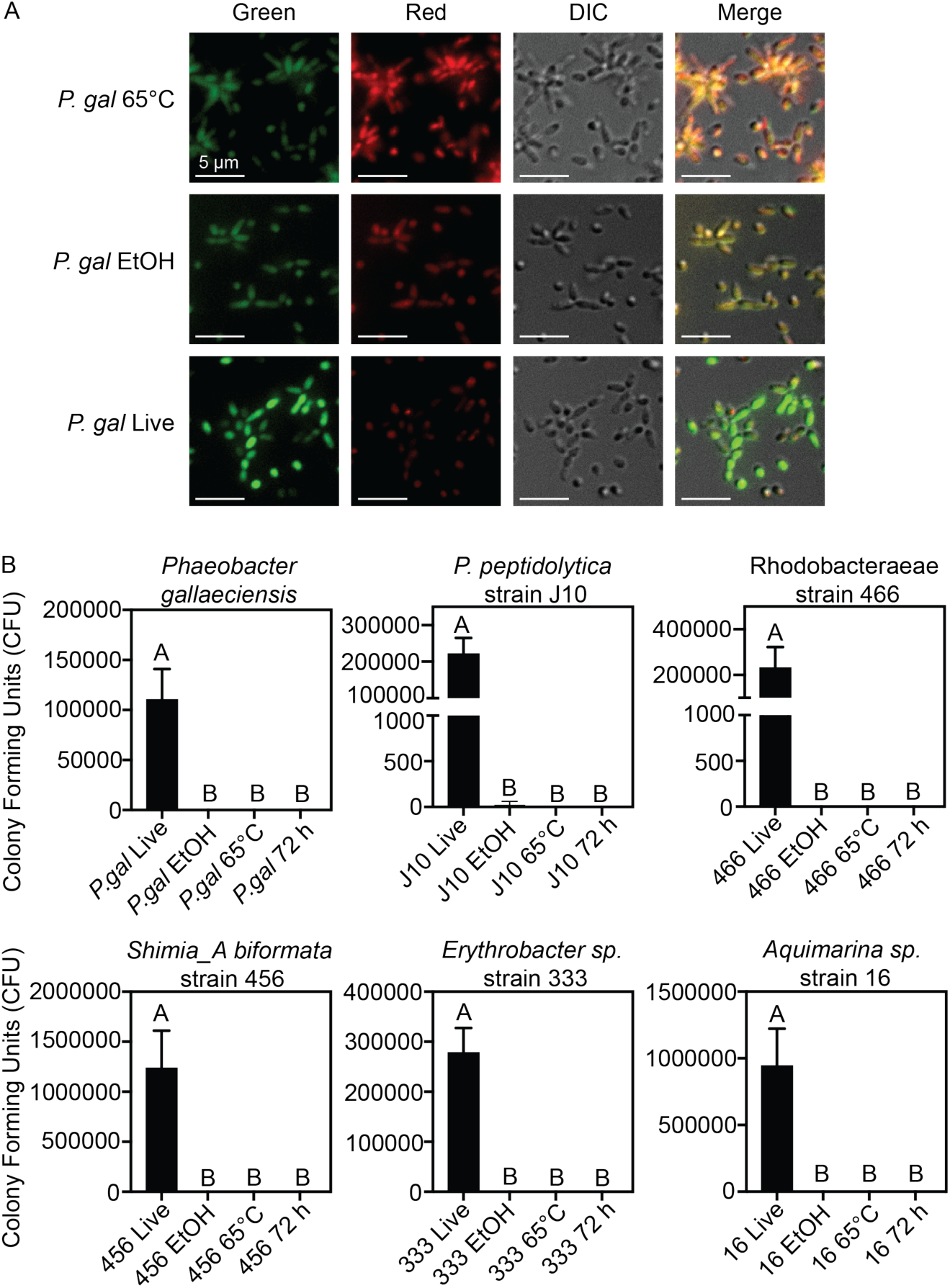
The ReefTiles method produces a non-living, bacteria-based surface coating. **(A)** Fluorescence and DIC overlay micrographs of *Phaeobacter gallaeciensis (P. gal)* biofilms after treatments: untreated (Live), 70% Ethanol (EtOH), dried at 65°C for 2 h. Live cells stained with Acridine Orange (Green) and dead cells stained with Propidium iodide (Red). **(B)** Bacterial colony forming units (CFU) of Rhodobacteraceae strain 466, *Shimia_A biformata* strain 456, *Pseudoalteromonas peptidolytica* strain J10, *Erythrobacter sp.* strain 333, *Aquimarina sp.* strain 16, and *Phaeobacter gallaeciensis* biofilms after treatments: untreated (live), 70% Ethanol, dried at 65°C for 2 h or dried at ambient temperature for 72 h. Data represents the average of three technical replicates. Error bars represent standard deviation (STD). One-way ANOVA (***p<0.0005) with letters representing Tukey’s multiple comparisons test.

### ReefTiles stimulate settlement of tubeworm larvae

Having established that the ReefTile coating is stable and non-viable, we next tested whether the inactivated bacterial films retained biological activity that could stimulate larval settlement including attachment to the surface. Using the model biofouling tubeworm *Hydroides elegans*, we compared the inductive capacity of ReefTiles prepared with a species from the Roseobacter clade *Phaeobacter gallaeciensis* to that of conventional live *P. gallaeciensis* bacterial biofilms and uncoated control tiles. Across multiple trials, ReefTiles strongly induced metamorphosis, demonstrating that the stimulatory molecules remain active after drying (**Fig. 3A-B**). Notably, ReefTiles produced settlement rates equivalent to or exceeding those of living biofilms, while blank tiles without bacterial coating elicited minimal response. These results indicate that the biochemical cues responsible for inducing settlement are heat- and desiccation-tolerant. To determine if induction by ReefTiles results in successful larval outcomes, we tested for tubeworm survival 1-week after settlement. Larvae that settled on ReefTiles survived better than living bacterial biofilms (**Fig. 3C**). To test the shelf-life of ReefTile coatings to determine their stability, we stored conditioned ReefTiles for 1-month at ambient temperature and then tested settlement. We observed similar levels of larval settlement on 1-month old ReefTiles (33%) when compared to fresh ReefTiles (43%) (**Fig. 3D**).

**Fig 3.**
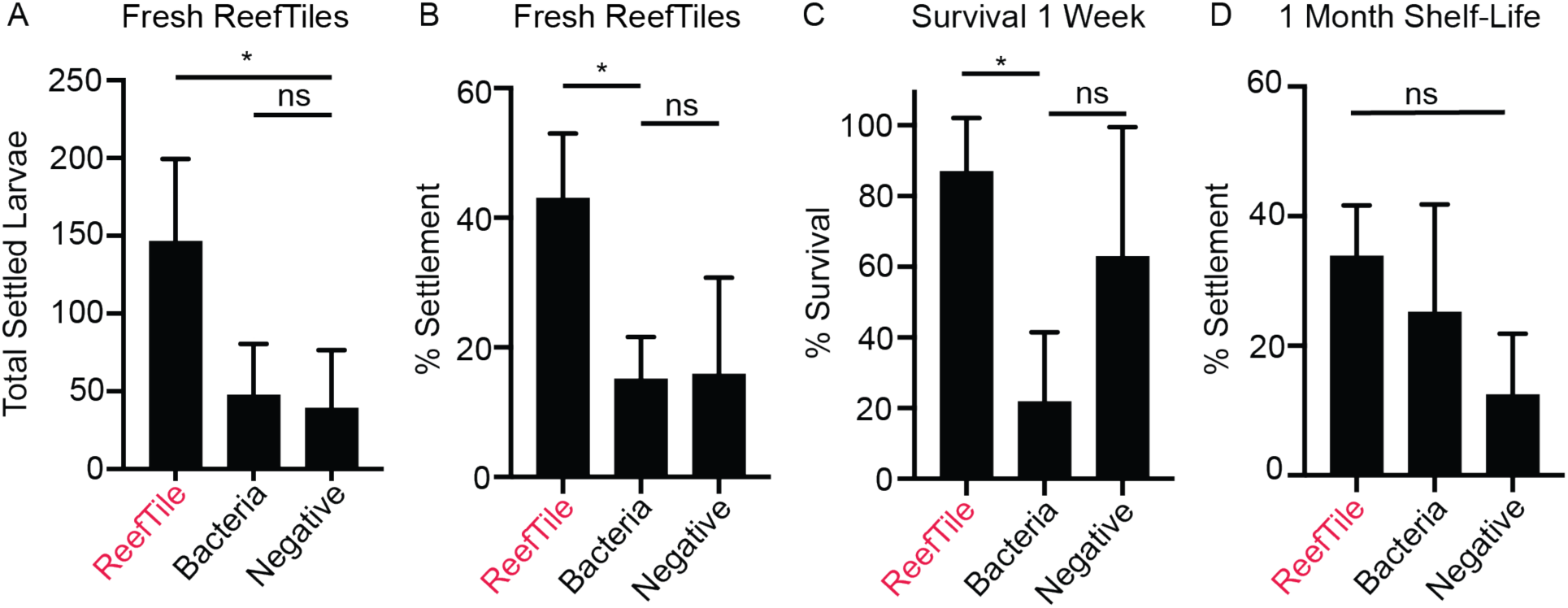
ReefTile coating promotes the settlement and survival of tubeworm larvae. **(A)** Total settled larvae (approximately 150 larvae per treatment with 3 biological replicates n=3) exposed to a biofilm of *P. gallaeciensis* ReefTiles, living bacterial biofilm (Bacteria) and tiles conditioned in FSW without bacterial coating (Negative). Error bars represent standard deviation. One-way ANOVA with Tukey’s multiple comparisons test (*p=0.0467). **(B)** Percent settlement of larvae (approximately 150 larvae per treatment with 3 biological replicates n=3) exposed to a biofilm of *P. gallaeciensis* ReefTiles, living bacterial biofilm (Bacteria) and tiles conditioned in FSW without bacterial coating (Negative). Error bars represent standard deviation. One-way ANOVA with Tukey’s multiple comparisons test (*p=0.0478). **(C)** Percent survival of larvae (approximately 150 larvae per treatment with 3 biological replicates n=3) exposed to a biofilm of *P. gallaeciensis* ReefTiles, living bacterial biofilm (Bacteria) and tiles conditioned in FSW without bacterial coating (Negative) after 1-week. Error bars represent standard deviation. One-way ANOVA with Tukey’s multiple comparisons test (*p=0.0143). **(D)** Percent settlement of larvae (approximately 150 larvae per treatment with 3 biological replicates n=3) exposed to a biofilm of *P. gallaeciensis* ReefTiles, living bacterial biofilm (Bacteria) and FSW without bacterial coating (Negative), conditioned 1-month prior. Error bars represent standard deviation. One-way ANOVA with Tukey’s multiple comparisons test (ns).

Together, these experiments demonstrate that ReefTiles preserve the inductive activity of bacteria while eliminating the need for living biofilms. The coating therefore provides a standardized, reproducible surface that can reliably trigger metamorphosis of tubeworm larvae, establishing a foundation for broader application across marine invertebrate species.

### Strain-specific bacterial cues are retained in ReefTile coatings for tubeworm and coral larvae

To identify effective bacterial species for ReefTile applications, we used the panel of six bacterial strains with the ReefTiles method (**Table S1**) (Alker et al. 2023a), and tested their ability to induce *Hydroides elegans* (tubeworm) metamorphosis. Tubeworm larvae settled in response to some strains shown to induce coral settlement (Rhodobacteraceae strain 466, *S. biformata* strain 456, and *Erythrobacter sp*. strain 333) with most ReefTile dried strains resulting in similar or greater levels of settlement than live biofilms (**Fig. 4A**). Some coral-inducing isolates such as *Pseudalteromonas peptidolytica* strain J10 and *Aquimarina sp.* strain 16 did not induce tubeworm settlement, demonstrating that the ReefTile dried biofilm preserves inductive activity and does not create it.

**Fig 4.**
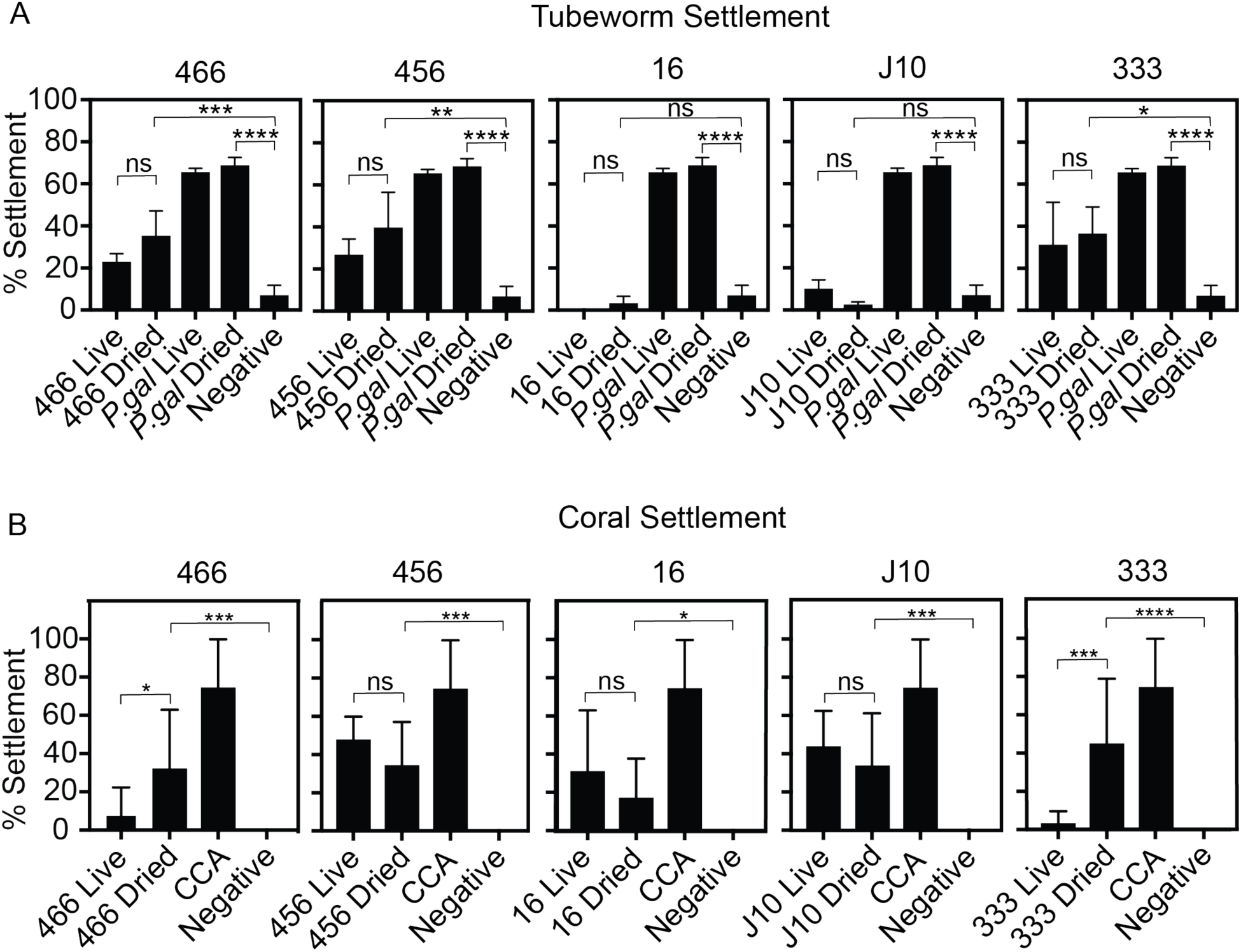
ReefTile coating of multiple bacterial species stimulates settlement of tubeworm and coral larvae. **(A)** Percent settlement of *Hydroides elegans* larvae (approximately 150 larvae per treatment with 3 biological replicates n=3) exposed to a live or ReefTile dried biofilm of Rhodobacteraceae strain 466, *Shimia_A biformata* strain 456, *Pseudoalteromonas peptidolytica* strain J10, *Erythrobacter sp.* strain 333, *Aquimarina sp.* strain 16, and *Phaeobacter gallaeciensis*. Settlement tiles conditioned in FSW without bacterial coating were used as negative control while *Phaeobacter gallaeciensis* live biofilm was used as positive control. Error bars represent standard deviation. One-way ANOVA with Tukey’s multiple comparisons test (*p<0.05, **p<0.005, ***p<0.0005, ****p<0.0001). **(B)** Percent settlement of *Acropora kenti* larvae (6 larvae per treatment with 6 biological replicates n = 6) exposed to a live or ReefTile dried biofilm of Rhodobacteraceae strain 466, *S. biformata* strain 456, *Pseudoalteromonas peptidolytica* strain J10, *Erythrobacter sp.* strain 333, and *Aquimarina sp.* strain 16. Settlement tiles conditioned in FSW without the bacterial coating were used as the negative control while crustose coralline algae (CCA) was used as a positive control. Error bars represent standard deviation. Fisher’s Exact Test (*p<0.05, **p<0.005, ***p<0.0005, ****p<0.0001).

To determine whether ReefTiles could enhance larval settlement in reef-building corals, we conducted assays using *Acropora kenti* larvae and five inductive bacterial strains previously shown to promote coral larval settlement (**Table S1**), including *Pseudalteromonas peptidolytica* strain J10 (Tebben et al. 2011) and four strains isolated from inductive biofilms (Rix et al., 2024). Biofilms of live bacterial strains and dried ReefTiles were compared for their ability to induce settlement with attachment. Data was analyzed as raw counts of larvae using Fisher’s Exact Test, while data displayed is the averaged percent settlement across biological replicates. Larvae exposed to ReefTiles exhibited significantly higher rates of settlement and normal polyp formation than those exposed to uncoated control tiles for all strains tested (**Fig. 4B; Fig. S1**). In two strains (*Erythrobacter sp.* strain 333 and Rhodobacterceae strain 466), settlement frequencies on ReefTiles were higher compared to those observed on live biofilms, while the three remaining strains (*Pseudoalteromonas peptidolytica* strain J10, *S. biformata* strain 456 and *Aquimarina sp.* strain 16) showed equivalent settlement rates between the ReefTile and live biofilms. In some cases, metamorphosis was induced without attachment (**Fig. S1**). This was particularly prevalent in the *Pseudoalteromonas peptidolytica* strain J10, where all larvae were either settled or underwent metamorphosis in both the ReefTile and live biofilms. Taken together, these results indicate that the drying process preserved the inductive biomolecules responsible for coral larval settlement.

Given the strong inductive properties of the *Pseudoaltermonas peptidolytica* strain J10 isolate (100% larvae either settled or metamorphosed), we tested the *Pseudoaltermonas peptidolytica* strain J10 ReefTile on an additional four coral species (**Fig. S2; Table S2**). In three of the four species, the ReefTile induced equivalent settlement rates to live biofilms, with almost 100% of larvae either settled or metamorphosed in the two acroporids, and over 50% of larvae either settled or metamorphosed in *Goniastrea retiformis*. Although nearly 50% of *Echinophyllia aspera* larvae were either settled or metamorphosed in response to the ReefTile, this was close to 75% in response to live biofilms. Overall, ReefTiles induced settlement in all five coral species tested, including both acroporid and non-acroporid species from the families Acroporiidae, Meruliniidae and LobophyllidaeThis establishes the proof-of-concept that ReefTiles can be customized using specific inductive bacterial taxa to promote coral recruitment in restoration settings.

## DISCUSSION

Our results demonstrate that a simple, non-living bacterial coating can reproducibly stimulate metamorphosis and attachment of marine invertebrate larvae, addressing a persistent challenge in restoration and aquaculture systems where larval settlement is often inconsistent. By drying and inactivating inductive bacteria while preserving their stimulatory biomolecules, the ReefTiles approach translates microbial cues into a stable and easy-to-use coating that can be applied across different substrate types. This coating method represents a practical advance in applying stimulatory microbes to promote settlement while remaining environmentally safe and readily scalable.

A specific advantage of the ReefTiles coating method is its ability to be coupled with existing innovations in substrate composition that influence larval settlement, creating opportunities for synergistic enhancement of metamorphosis and attachment. In this study, we focused on ceramic or concrete tiles as traditional restoration surfaces for a consistent model surface, but future work will explore how substrate composition can be tuned in concert with bacterial species coatings to further enhance larval settlement and post-metamorphic success. Substrate mineralogy and chemistry are known to shape settlement responses, with materials enriched in calcium carbonate, strontianite, or silica promoting recruitment through the release of ions and the formation of favorable surface textures (Levenstein et al., 2022; Maboloc et al., 2025). These same substrate properties also modulate microbial colonization, fostering biofilms and crustose coralline algae (CCA) that naturally stimulate metamorphosis (Lei et al., 2021; Rippen et al., 2025). Integrating the ReefTiles microbial coating with such optimized substrates could therefore amplify settlement cues and extend applicability across a wider range of materials.

In this study, we demonstrated that coatings made from specific inductive bacteria—*Phaeobacter gallaeciensis*, Rhodobacteraceae, *Shimia_A biformata*, *Aquimarina sp.*, *Erythrobacter sp*., *Pseudoalteromonas peptidolytica*—can effectively stimulate larval settlement and metamorphosis in both tubeworms and corals. We selected Gram-negative, non–spore-forming bacteria from the Alphaproteobacteria and *Pseudoalteromonas* lineages because these groups have previously been shown to stimulate metamorphosis in diverse marine invertebrates (Alker et al., 2020; Alker, Farrell, Aspiras, et al., 2023; Alker, Farrell, Demko, et al., 2023; Sneed et al., 2014b). Many other bacterial strains have been demonstrated to induce settlement in a range of invertebrates, and could be adapted for use in the ReefTiles method (Negri et al., 2001; Petersen et al., 2021). While our current approach focuses on single-strain coatings, mixed-species coatings could further improve the robustness and breadth of larval responses by combining complementary bacterial products, mimicking the complexity of natural biofilms (O’Brien et al., 2026; Petersen et al., 2021). Other groups have explored similar microbial interfaces, including living hydrogel coatings that maintain active bacteria (Levy et al., 2025), CCA-associated bacterial consortia that act synergistically with algal metabolites to induce coral settlement (Gómez-Lemos et al., 2018; Siboni et al., 2020), and chemical inducers embedded in agar hydrogels or biomaterials (Briggs et al., 2026; Kundu et al., 2025). Together, these studies provide a foundation for developing next-generation microbial coatings that integrate multiple inductive species and biochemical classes, expanding the versatility and ecological relevance of the ReefTiles technology.

In summary, ReefTiles represent a versatile and practical advance in applying microbial signaling to marine restoration and aquaculture. By stabilizing bacterial settlement cues within an inert coating, this technology bridges the gap between mechanistic understanding of metamorphosis and scalable, field-ready application. The ability to preserve bioactivity without maintaining living cells offers distinct advantages in biosafety, reproducibility, and long-term storage. Importantly, the method can be adapted to different substrates and bacterial species, allowing tailored combinations for target organisms or environments. As a modular platform, ReefTiles can be combined with other emerging restoration technologies, including mineral-engineered substrates and settlement-inducing bacterial strains from CCA, enabling future efforts to pair stimulatory bacteria with optimized substrate properties. Collectively, this approach establishes a foundation for developing next-generation bioactive materials that use natural microbial chemistry to direct settlement, growth, and ecological restoration in the sea.

## MATERIALS AND METHODS

### Bacterial strains and culture methods

Marine bacteria used in this study are found in Table S1, strains were cultured on Marine Broth (MB) 2216 (BD Difco). Marine bacteria were incubated at 25°C, and cultures were shaken at 200 rpm. All liquid cultures were inoculated with a single colony and incubated between 16 and 18h.

### ReefTile coating method

Marine bacteria were inoculated into MB broth and grown at 25°C shaken at 200 rpm for 16 to 18 h. Bacterial cultures were washed 3 times with 0.22 µm filtered artificial sea water (ASW). Optical density was quantified at 600 nm (OD 600) and cultures were diluted to an OD of 1. Settlement tiles were soaked in deionized water (DI) as recommended, on average 1-3 days with water changes every day. Settlement tiles both ceramic (ReefH20 Frag Discs, Arizona USA) and concrete (made in-house with custom molds and Dunlop Concrete Resurfacer) were soaked in OD 1 bacteria for 1 h. Settlement tiles were dried at room temperature <72 h or heated to 65°C for 2 hours. After drying the tiles were stored at room temperature until use.

### Live Dead Staining and Bacterial Counts

Rhodobacteraceae strain 466, *Shimia_A biformata* strain 456, *Pseudoalteromonas peptidolytica* strain J10, *Erythrobacter sp.* strain 333, *Aquimarina sp.* strain 16, and *Phaeobacter gallaeciensis* were struck onto Marine Broth 2216 (BD Difco) agar plates; a single colony was inoculated into MB liquid culture and grown for 16-18 hours at 25℃. Bacterial cultures were washed 3 times with 0.22 µm filtered artificial sea water (ASW). Bacterial cultures were quantified by optical density at 600 nm (OD 600) and were diluted to an OD of 1. Biofilms were seeded onto 35 mm petri dishes with 14 mm microwell coverglass (MatTek Life Sciences, Ashland, MA) for imaging or 96-well cell culture plates for bacterial counts. Biofilms formed for 1 h and then washed unbound bacteria with 1x phosphate buffered saline (PBS). For live biofilms staining was added in the working solution 5 µg/mL of Acridine Orange (AO) and 10 µg/mL of Propidium iodide (PI) in 1x PBS. AO is able to enter both live and dead cells, in live cells the signal is green. When PI enters dead cells the signal of AO is partially or completely absorbed into the PI signal due to the FRET phenomenon, leading to predominately or only red PI signal in dead cells. Stain was incubated in the dark for 30 mins and then washed with 1x PBS. Ethanol treated biofilms had 70% Ethanol added and incubated at room temperature for 30 mins before washing with 1x PBS and applying stain. Heat treated biofilms were incubated at 65℃ for 2 h and then stain was applied. Stain remained on samples for 30 mins and then washed with 1x PBS.

Bacterial biofilms in the 96-well culture plate were treated the same as imaging dishes with the exception of staining. Instead triplicate wells were resuspended in 100 µL of MB broth and plated onto a MB agar plate for each technical replicate incubating at 25℃ overnight. Live biofilms were diluted to either 1:500 or 1:1000 through serial dilution depending on strain growth rate and then plated on MB agar plates. Bacterial colonies were counted after overnight growth, the average was calculated from the counts of three technical replicate plates for each condition. Live biofilm conditions were calculated based on dilution factor for each technical replicate plate.

### Microscopy

Microscopy was performed using a Zeiss Axio Observer.Z1 inverted microscope equipped with an Axiocam 820 mono camera and Plan-Apochromat 100×/1.4 DICIII objective. The Zeiss eGFP filter set 38 HE (000000-1031-346) was used to capture Acridine Orange staining and the Zeiss mRFP filter set 63 HE (489063-0000-000) was used to capture Propidium iodide staining. Images were captured using the same fluorescence exposure times for all samples. ZEN 2 software was used for image processing using identical fluorescence gating and mild unsharp masking across all conditions.

### Tubeworm settlement assays

Biofilm settlement assays were conducted as previously described (Huang et al. 2012; Shikuma et al. 2014). Briefly, marine bacteria were struck onto Marine Broth 2216 (BD Difco) agar plates. A single colony was inoculated into MB liquid culture and grown for 16 to18 hours at 25℃. Bacterial cultures were washed 3 times with 0.22 µm filtered artificial sea water (ASW). Bacterial cultures were quantified by optical density at 600 nm (OD 600) and were diluted to an OD of 1. To form the biofilm 100 µL of diluted bacterial culture was seeded into a 96-well cell culture plate. ASW negative control wells had 100 µL of ASW added to the well. The biofilm formed for 1 hour and was then washed with ASW 2 times. *Hydroides elegans* competent larvae (Day 6-7) were added to each well at a density of 50 larvae per well with 3 technical replicate wells. Settlement data was analyzed as percentages to accommodate slight variances in the number of tubeworm larvae across biological replicates.

### Coral settlement assays

Bacterial strains were revived from frozen glycerol stocks through inoculation (∼10 μL) onto ½ MB agar and sub-cultured onto fresh media as required to maintain viability throughout the experiment (1–2 weeks). For biofilm development, bacterial isolates were grown for 24–48 h in filtered ½MB at 28°C and 180 RPM as required to reach sufficient visible cell density. Bacterial cultures were washed two times with sterile 0.22 µm filtered seawater (FSW) to remove growth media. Cell densities were quantified by optical density at 600 nm (OD 600) and diluted or concentrated as required to reach an OD of 1. Concrete settlement tiles (8 × 8 × 5 mm^3^) were prepared and soaked in FSW for one week to leach out any concrete-associated chemicals and then sterilised by autoclave. For each strain, biofilms were formed on the tiles by placing them in sterile petri dishes filled with the resuspended bacterial culture. Negative control tiles were prepared in sterile FSW. The tiles were incubated in the dark at 28°C for 24 h to allow biofilms to develop and then washed in sterile FSW. Live biofilms were immediately used in settlement assays. ReefTiles were dried at room temperature for 24 h and stored at room temperature until use (within 1–2 weeks).

Coral collection, spawning, larval culture, and settlement assays were conducted as previously described (Abdul Wahab et al., 2023; O’Brien et al., 2025). Briefly, colonies from five species of coral (*Acropora anthocercis, A. kenti, Acropora* sp. nov. aff. *kenti* [formerly *A. tenuis* (Bridge et al., 2024)], *Echinophyllia aspera, Goniastrea retiformis*) were collected from the Palm Island Group (18°45’56.4”S 146°32’2.58”E) approximately one week before the predicted 2024 Great Barrier Reef spawning events following the October and November full moons and brought to the National Sea Simulator (SeaSim) at the Australian Institute of Marine Science (AIMS). Colonies were held in outdoor semi-recirculating aquaria and isolated into 60 L tanks for gamete collection. Gametes from all parent colonies of the same species were pooled for cross fertilisation and rinsed embryos were transferred to 70 L flow-through tanks where larval cultures were maintained until used in settlement assays. Larval competency was assessed using crustose coralline algae (CCA) as a settlement inducer to ensure larvae were competent prior to being used in settlement assays.

Settlement assays were performed in sterile tissue culture 6-well plates (Corning, United States) with six replicate wells per bacterial strain per coral species. Each well contained 9 mL of FSW, six larvae, and a settlement tile containing a biofilm. Separate plates were run with CCA as a positive control, tiles incubated in FSW as a negative control, and blanks containing FSW with no settlement tiles (6 replicate wells each). Plates were incubated for 48 h at 28°C with a 12-hour light:dark cycle under lights providing 40 µmol⋅m^-2^⋅s^-1^ (Aqualina, Australia). After 48 h, larval settlement in each well was scored using a stereo microscope with larval responses grouped into five categories: (i) firmly attached without metamorphosis (attached), (ii) metamorphosis without attachment (metamorphosis), (iii) firmly attached and metamorphosed (settled), (iv) disintegrating or other signs of mortality (dead), and (v) swimming or stationary but no attachment (not settled).

### Statistical analysis of coral larval settlement

To determine whether settlement differed between the ReefTile and live bacteria treatments, a Fisher’s exact test was performed on a 2 x 2 contingency table for each isolate. Similarly, a Fisher’s exact test on a 2 x 2 contingency table for each isolate was used to evaluate whether settlement differed between the ReefTiles treatment and negative controls. Larval responses were grouped as settled or not settled (including categories i, ii, iv and v above), with total counts summed across replicates (n=6). Odds ratios were calculated as a measure of effect size, and a Benjamini-Hochberg correction was applied to account for multiple comparisons across all tests within a coral species. . All analyses were conducted in R (v4.4; (R Core Team 2024)) using base functions and the tidyverse package (v2.0) (Wickham et al., 2019).

## AUTHOR CONTRIBUTIONS

**M.V.F.:** Conceptualization; Methodology; Investigation; Formal analysis; Data curation; Writing – original draft; Writing – review & editing; Supervision; Funding acquisition. **L.R.:** Methodology; Investigation; Formal analysis; Data curation; Writing – original draft; Writing – review & editing; Funding acquisition. **P.A.O.:** Methodology; Investigation; Formal analysis; Data curation; Writing – original draft; Writing – review & editing. **T.L.D.:** Investigation; Writing – review & editing. **S.M.:** Investigation; Writing – review & editing. **F.K.:** Investigation; Resources. **N.J.S.:** Conceptualization; Methodology; Investigation; Formal analysis; Data curation; Writing – original draft; Writing – review & editing; Resources; Supervision; Funding acquisition.

## ACKNOWLEDGEMENTS

The authors acknowledge the Munbarra People, the traditional custodians of the land and sea country from which the corals used in this study were collected, and we thank them for their permission to use the larvae produced from these corals. We are grateful to the staff at the National Sea Simulator (SeaSim) and Coral Aquaculture and Deployment team (RRAP), with special thanks to Muhammad Abdul Wahab, for assistance in spawning corals and larval rearing. The authors acknowledge the Kumeyaay people, for millennia, the Kumeyaay people have been a part of this land. This land has nourished, healed, protected and embraced them for many generations in a relationship of balance and harmony. As members of the San Diego State community we acknowledge this legacy. We promote this balance and harmony. We find inspiration from this land; the land of the Kumeyaay

## SUPPLEMENTARY MATERIALS

**Table S1:** Strains used in this study.

| Strain no. | Strain | Identifier | Genotype | Source |
| --- | --- | --- | --- | --- |
| NJS412 | *Phaeobacter gallaeciensis*<br>ATCC 700781 (DSM 26640) | *P. gal* | StR | (Ruiz-Ponte et al., 1998) |
| NJS893 | Rhodobacteraceae strain 466 | 466 | Wild-type | (Rix et al., 2024) |
| NJS894 | *Shimia_A biformata* strain 456 | 456 | Wild-type | (Rix et al., 2024) |
| NJS895 | *Aquimarina* sp. strain 16 | 16 | Wild-type | (Rix et al., 2024) |
| NJS896 | *Pseudoalteromonas*<br>*peptidolytica* strain J10 | J10 | Wild-type | (Tebben et al., 2011) |
| NJS897 | *Erythrobacter* sp. strain 333 | 333 | Wild-type | (Rix et al., 2024) |

**Table S2:** Supporting statistics for Figure S2.

| **Coral Species** | **Isolate** | **Comparison** | **Odds Ratio** | **p_value** | **p_adjusted** | **Significance** |
| --- | --- | --- | --- | --- | --- | --- |
| *A. anthocersis* | J10 | Dry vs Live | 1.266 | 0.775 | 0.886 | FALSE |
| *Acropora sp.* | J10 | Dry vs Live | 1.374 | 0.770 | 0.886 | FALSE |
| *E. aspera* | J10 | Dry vs Live | 0.250 | 0.019 | 0.039 | TRUE |
| *G. retiformis* | J10 | Dry vs Live | 0.892 | 1.000 | 1.000 | FALSE |
| *A. anthocersis* | J10 | Dry vs Neg | 0.781 | 0.774 | 0.886 | FALSE |
| *Acropora sp.* | J10 | Dry vs Neg | Inf | 0.001 | 0.006 | TRUE |
| *E. aspera* | J10 | Dry vs Neg | Inf | 0.008 | 0.022 | TRUE |
| *G. retiformis* | J10 | Dry vs Neg | 21.456 | 0.000 | 0.002 | TRUE |

**Figure S1.**
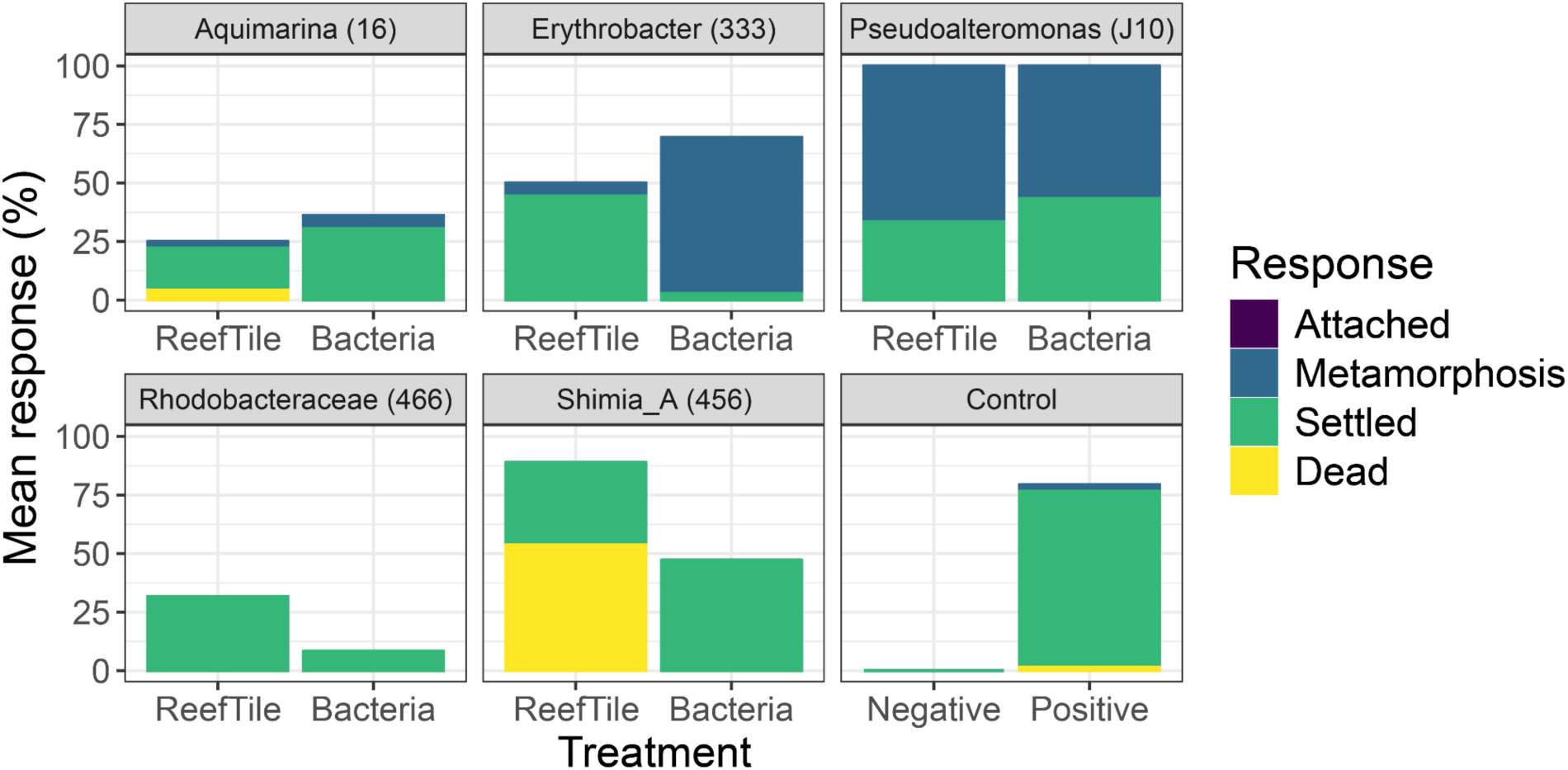
Mean settlement responses of *Acropora kenti* larvae (6 larvae per treatment with 6 biological replicates n = 6) exposed to ReefTiles and live biofilms. Each panel represents a different bacterial isolate. Settlement tiles without the bacterial coating were used as the negative control while crustose coralline algae was used as a positive control. Attached refers to larvae firmly attached without metamorphosis. Metamorphosis refers to larvae that underwent metamorphosis without attachment.

**Figure S2.**
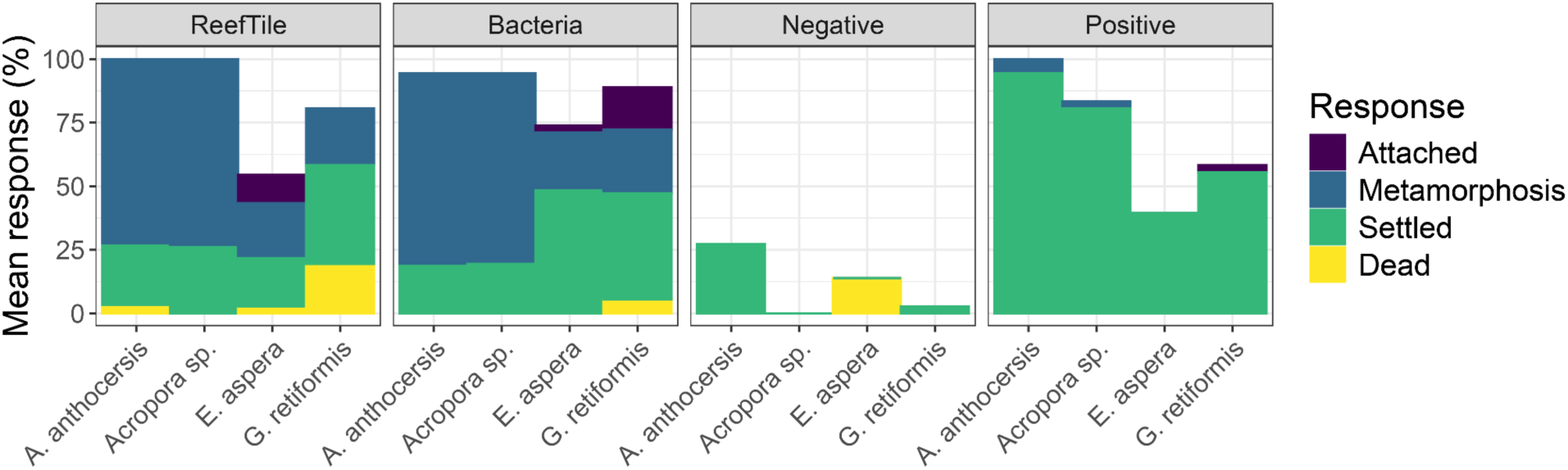
Mean settlement responses of larvae from an additional four coral species (6 larvae per treatment with 6 biological replicates n = 6) exposed to ReefTiles and live biofilms of *Pseudoalteromonas peptidolytica* strain J10. Settlement tiles without the bacterial coating were used as the negative control while crustose coralline algae was used as a positive control. Attached refers to larvae firmly attached without metamorphosis. Metamorphosis refers to larvae that underwent metamorphosis without attachment.

